# Overexpression of HOP2 induces developmental defects and compromises growth in Arabidopsis

**DOI:** 10.1101/2021.08.17.456113

**Authors:** Ameth N. Garrido, Therese Francom, Sakina Divan, Mohamad Kesserwan, Jenya Daradur, C. Daniel Riggs

## Abstract

HOMOLOGOUS PAIRING 2 (HOP2) is a predominantly meiotic protein that plays a pivotal role in homologous chromosome pairing in organisms as diverse as yeast and mammals. While generating HOP2::GFP reporter lines, we identified two Arabidopsis T-DNA insertion mutants, *stunted1* (*std1*) and *stunted2* (*std2*) that exhibit pleiotropic phenotypes, including fasciated stems, altered phyllotaxy, floral organ defects, reduced fecundity, and an overall reduction in growth properties. TAIL-PCR followed by sequencing revealed several insertions near genes, but genotyping showed that none of the insertions are causal. Analysis the *std* mutants by qRT-PCR, and analysis of dexamethasone inducible HOP2 transgenic plants demonstrated that the *std* phenotypes are associated with ectopic/overexpression of HOP2. Based on the postulated mechanisms of HOP2 action, we speculate on how overexpression leads to these developmental/growth defects.

## Introduction

Meiosis is an evolutionarily conserved cell division mechanism that generates genetic diversity in the meiotic products, either gametes or spores, depending on the organism. After cells are committed to meiosis, the duplicated chromosomes undergo intricate movements and interactions that promote the alignment of homologous pairs, which subsequently exchange segments during recombination. These movements may involve the attachment of telomeres to either the nuclear membrane or the nucleolus, the clustering of centromeres, and the synapsis of the chromosome arms (Unhavaithaya and Orr-Weaver, 2013; Da Ines and White, 2015; Hurel et al., 2018; Sepsi and Schwarzacher, 2020; Tian et al., 2020). As synapsis proceeds, a proteinaceous matrix, the synaptonemal complex (SC) is assembled between the paired homologs and commitment to crossing over occurs. Double strand breaks (DSBs) are induced during meiosis and they are repaired using Homologous Recombination repair (HR). Many eukaryotes, including Arabidopsis, rely on HR repair of DSBs to accomplish both synapsis and crossing over. In most organisms the concerted action of numerous proteins, including RAD51, DMC1, MND1 and HOP2 are required for these processes (reviewed by Ranjha et al., 2018; Crickard and Greene, 2018). RAD51 and DMC1 are both RecA type recombinases that promote the formation of the ssDNA nucleofilaments that invade dsDNA and initiate strand exchange that can lead to a crossover. HOP2 and MND1 are accessory proteins that form a heterodimer, and are proposed to tether the ssDNA to dsDNA, providing a critical level of stabilization. In yeast and mammals, null mutations in HOP2 or MND1 lead to a general failure in homologous chromosome SC formation and repair of DSBs, resulting in meiotic arrest (Leu et al., 1998; Tsubouchi and Roeder, 2003; Petukova *et al*. 2003). In Arabidopsis and rice, HOP2 and MND1 null mutants also have a general failure of SC formation, but DSBs are repaired and there is no meiotic arrest (Kerzendorfer et al., 2006; Panoli et al., 2006; Stronghill et al., 2010, Shi et al., 2019). Despite the general lack of SC formation in Arabidopsis *hop2* or *mnd1* mutants, aberrant chromosome segregation with chromatin bridges and fragments was observed, suggesting that HOP2 and/or MND1 have a dual role both in promoting SC formation between homologs or preventing stable associations from forming between nonhomologous chromosomes (Farahani-Tafreshi et al., 2021). Mechanistically, HOP2 and MND1’s action is not completely understood, but *in vitro* studies provide evidence that the HOP2/MND1 heterodimer modulates the activity of the RAD51 and DMC1 recombinases and HOP2, in the absence of MND1 is a potent recombinase (Petukova et al., 2005; Neale and Keeney 2006; Ploquin et al., 2007; Bugreev et al., 2014; Tsubouchi et al., 2020).

To support our on-going work on Arabidopsis HOP2, we attempted to generate a reporter gene transgenic lines by linking the HOP2 promoter and coding region to a construct encoding the fluorescent reporter GFP. We were successful in generating reporter lines, which had a normal meiosis and expressed GFP in the meiotic cells, but we also observed some transformants that displayed a range of developmental abnormalities. Here we report the characterization of two such mutants that we termed *stunted1* (*std1*) and *stunted2* (*std2*). We propose that ectopic/overexpression of HOP2 is responsible for the *std* phenotypes.

## Experimental Procedures

The *hop2-1* mutant was obtained from the Arabidopsis Biological Resource Center (ABRC). Plants were grown in Conviron growth chambers on a 16hr day/8hr night regimen under fluorescent lights (~125μE/m^2^) in Premier Promix PGX medium.

### Generation of std mutants

PCR was used to amplify the wildtype HOP2 gene to generate a construct containing 4681bp preceding the start codon, and creating an in-frame fusion with the coding region of GFP in a pEGAD derivative. Primer sequences and the construction strategy are given in Table S1. DNA sequencing confirmed the translational in-frame fusion. Agrobacterium mediated transformation of either Columbia or Landsberg erecta (L*er*) plants was employed and transformants selected by growth on 0.5X MS plates containing 10ug/ml BASTA.

### Microscopy

Light microscopy and image capture were undertaken by using a Leica MZ 7.5 or Nikon SMZ1500 dissecting microscopes. For scanning electron microscopy, samples were prepared and examined by employing a Hitachi S-530 scanning electron microscope at 25 kV, as described by Douglas et al. (2002), and images were captured using the Quartz PCI version 8 system.

### TAIL-PCR

Genomic DNA from *std1* and *std2* mutants was prepared and used in TAIL-PCR essentially as described by Sessions and coworkers (2002). Two rounds of amplification were conducted using New England Biolabs Q5 polymerase. In the primary reactions a T-DNA border primer was used with a combination of degenerate primers (see Table S1), and PCR parameters were as follows: 94°C, 2min; 5 cycles of 98°C, 15 sec/62°C, 60 sec/72°C, 120sec; then 2 cycles of 98°C, 15 sec/25°C, 180 sec, with ramp rate 0.5°C per sec/72°C, 120sec; then 15 cycles of 98°C, 15 sec/68°C, 60 sec/72°C, 120sec/98°C, 15 sec/68°C, 60 sec/72°C, 120sec/98°C, 20 sec/44°C, 60 sec/72°C, 120sec/72°C, 180 sec. One microliter of the primary reaction was used in secondary reactions with parameters: 94°C, 2.5min; 5 cycles of 98°C, 20 sec/62°C, 60 sec/72°C, 120sec; then 15 cycles of 98°C, 15 sec/64°C, 60 sec/72°C, 120sec/98°C, 15 sec/64°C, 60 sec/72°C, 120sec/98°C, 20 sec/44°C, 60 sec/72°C, 120sec, then 5 cycles of 98°C, 20 sec/44°C, 60 sec/72°C, 120sec/, then 72°C, 180 sec. The reactions were subjected to agarose gel electrophoresis and DNA fragments visualized by staining with Red-Safe (Frogga Bio). The bands were resected and purified by employing a Bio-Basic ez10 gel purification kit. The purified DNA was then sequenced with the T-DNA border primer at the Hospital for Sick Children TCAG facility (Toronto).

### T-DNA insertion analysis

The status of T-DNA insertions into loci defined by TAIL-PCR sequencing was undertaken by the standard method of employing two PCR reactions. To test for an uninterrupted (wildtype) gene, primers that flank the T-DNA insertion site were used; to test for a T-DNA insertion, a gene specific primer/T-DNA border primer was employed. To examine the organization of the T-DNA insertions as tandem repeats, convergent or divergent insertions, we employed the relevant T-DNA border primers from one or both ends (see Table S1 for primers and strategy).

### DEX-inducibility construct and DEX assays

We generated a DEX inducible HOP2 construct by amplifying the HOP2 gene with primers that added attB sites (see Table S1), then mobilized the clone into pDONR207 by employing Gateway technology (Invitrogen) to generate an entry clone. The sequence was then inserted into pDEX to generate the destination clone that contains a C-terminal HA tag in frame with HOP2 (construct LC842). LC842 was mobilized into Agrobacterium GV3101 and used to transform L*er* plants by the floral dip method. Transgenic plants were selected on medium containing 0.5X MS salts, 5mM MES, pH5.7, 1% sucrose, 0.8% agar, and 10ug/ml BASTA.

For plate assays, LC842 and L*er* seeds were surface sterilized and germinated on either MS medium (L*er*) or MS medium containing BASTA (LC842), as outlined above. After 3 days of stratification at 4°C, plates were moved to room temperature under fluorescent lighting and allowed to grow for one week, then transferred to either control plates (MS medium) or MS plates containing 1uM dexamethasone (BioShop). Plates were imaged on day 0 (day of transfer) and day 5. For qRT-PCR experiments, seedlings were collected after 3 days and used to prepare RNA for analysis of transcript levels.

For *in terra* analyses, L*er* and LC842 seedlings were transferred from plates into pots containing Promix. Each 4-inch pot contained one seedling of each genotype. Control pots were spritzed with water containing 0.05% silwet, and DEX treatment was conducted by spritzing experimental pots with the same solution containing 1mM dexamethasone. Inflorescences were thoroughly wetted and spritzing was repeated every other day, for a total of four times. Plants were then imaged to capture phenotypic changes induced by DEX.

### qRT-PCR

RNA from plant tissues was purified and qRT-PCR conducted as described by Douglas et al., 2017. Primer and amplicon details are given in Table S1.

## Results

In the process of generating reporter lines to monitor *HOP2* expression, we identified a transformant that exhibited a variety of developmental abnormalities. At maturity, the plant was much shorter than wildtype and possessed thicker epinastic leaves, as well as thick and often fasciated stems (Figure 1). We also observed numerous instances of phyllotaxy defects, elevated levels of anthocyanins, and some instances of fused organs. Reproductive development was severely compromised. Floral buds appeared somewhat flattened and growth patterns of all organs were affected. Sepals and petals were shorter than wildtype and failed to enclose the reproductive whorls, creating an open flower phenotype. Stamens were disproportionally short and produced little or no pollen. The gynoecium was often open at the stigma and in some cases the carpels had failed to fuse. In general, very few stigmatic papillae were produced and fecundity was severely reduced. Given the overall slow growth and organ defects, we elected to call this mutant *stunted* (*std*). Occasionally the more normal looking pistils could be successfully fertilized with wildtype pollen and three general phenotypic classes were observed in the F2 progeny (Figure S1). The first class was nearly wildtype, but there were occasional minor phyllotaxy defects and fecundity issues. The second class of segregants was shorter than those of class1 and exhibited more extreme phenotypes, being overall very similar to the original *std* mutant. Class 3 segregants were severely stunted and the phenotypic defects observed in the original *std* mutant were exacerbated. Overall, there was broad range of defects that were sometimes not shared and the segregation ratio for the three classes was 2:2:1, which does not correspond to Mendelian ratios that point to recognized modes of inheritance.

**Figure 1:**
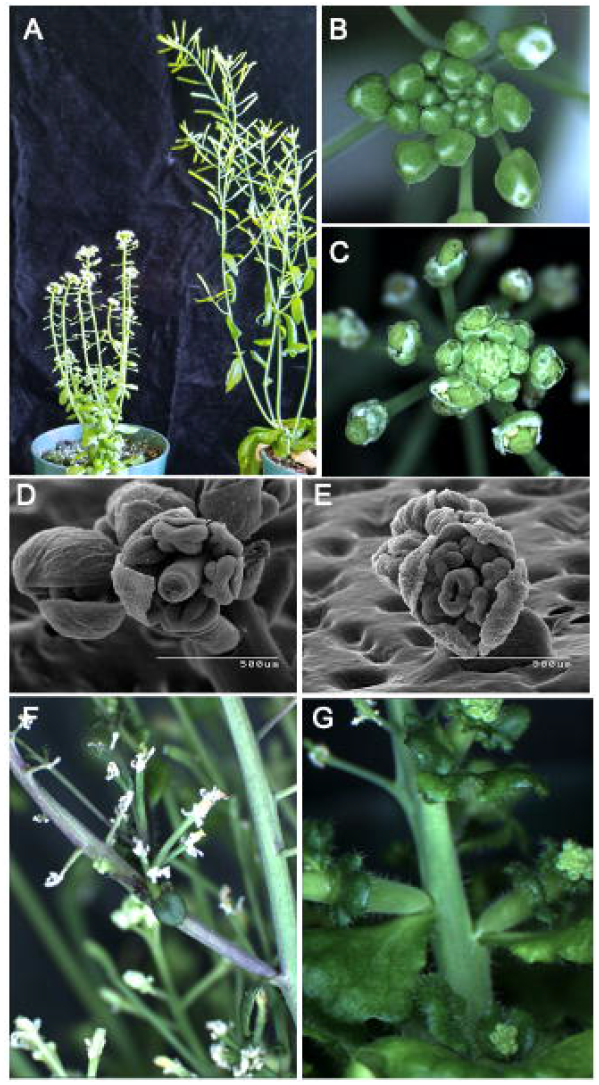
*std1* mutant phenotypes. A. An *std1* plant on the left and wildtype (L*er*) plant on the right, *std1* plant height is compromised and leaves are smaller, rounder and epinastic. B, C. inflorescences of wildtype (B) and *std1* (C), illustrating floral organ growth defects resulting in flattened and open buds. D, E. SEM photomicrographs illustrating sepal and pistil growth defects in wildtype (D) and *std1* (E). F. Phyllotaxy defects, reduced fecundity and elevated anthocyanin production in *std1*. G. Extreme phenotypes of *std* segregants that include thick and ruffled leaves, fasciated stems, and phyllotaxy and floral defects

### The stunted mutant is unlikely due to T-DNA insertion into a developmental regulatory gene

We hypothesized that *std1* arose due to random insertion of the reporter gene T-DNA into an important regulatory gene. To identify the insertion site, we performed thermal asymmetric interlaced PCR (TAIL-PCR) followed by sequencing of the products. In total, 38 sequences were obtained. About half of these sequences were revealed to be complex T-DNA rearrangements, head to tail tandem repeats of T-DNA, or were read-through sequences extending beyond the T-DNA left border sequence (Table 1). Sequence analysis of the transforming construct as well as the read-through insertions revealed no mutations within the LB and RB border sequences, and thus the mechanism(s) that produced these aberrant events is unknown, but the observations are not unprecedented (De Buck et al., 1999; De Neve et al., 1997; Wei et al., 2015). Amongst the TAIL-PCR sequences we found multiple hits for insertions in or near four genes: At4g16490, At3g21790, At3g47580, and At3g03460. In the first three cases, insertion occurred in the proximal promoter region of the genes and in the case of At3g03460, the insertion was 600bp downstream from the gene. To investigate whether disruptions in these genes is causal, we pollinated *std1* plants with wildtype pollen and genotyped F2 segregants for insertions in these four genes. Standard T-DNA genotyping was conducted for each set of plants (Figure S2). An analysis of two of the more extreme class 3 segregants and two intermediate phenotype plants (class 2) is shown in Table 2. The two class 3 extreme phenotype plants carried wildtype alleles for At3g47580 and At3g03460, eliminating these genes as being causal. Similarly, one of the two plants was wildtype for At4g16490, eliminating it as a candidate. The two class 2 plants were similar in that they were both mutant for At3g21790. However, some of the other intermediate phenotype plants were heterozygous for the insertion, discounting the involvement of At3g21790 (Figure S2). Moreover, it has been reported that insertions into the gene have no obvious phenotype (Dong et al., 2014), nor have any phenotypic defects been reported for insertions in the other three genes (At4g03460: Yu et al., 2020; others inferred from no reported phenotypes on TAIR). Thus, we found no apparent linkage between the *std1* mutant and insertions near the four genes. Because of the poor fecundity or possible lethality issues with *std1*, we eventually lost the line, but a subsequent transformation experiment gave rise to a phenotypically identical mutant that we termed *std2*. TAIL-PCR on *std2* DNA identified many similar instances of T-DNA rearrangements, LB/RB tandem repeats and LB read throughs, but also identified At3g03890 as having an insertion within the coding sequence (Table 3). This uncharacterized gene is predicted to encode a heme oxygenase. We examined the T-DNA status of At3g03890 in segregants of *std2*, and found that some mutant plants did not contain T-DNA in this gene, suggesting that disruption of At3g03890 is not causal (Figure S2). Thus, neither of the two independently isolated *std* mutants seems to possess insertions in common and none of the insertion sites identified by TAIL-PCR faithfully segregated with the mutant phenotype. Because many of the TAIL-PCR sequences were T-DNA repeats and rearrangements, this led us to re-examine the transforming construct and the expression of genes around the *HOP2* locus.

**Table 1:** TAIL-PCR of *std1* reveals multiple gene insertions and complex integration events

| Sequence count | Identity/annotation |
| --- | --- |
| 13 | Sequence reads through LB into vector DNA |
| 10 | Insertion upstream from At4g16490<br>• encodes an Armadillo repeat protein |
| 4 | T-DNA tandem repeat |
| 3 | Complex T-DNA rearrangement or PCR artifact |
| 3 | Insertion upstream from At3g21790<br>• encodes UDP-glucosyltransferase 71B7 |
| 3 | Insertion upstream from At3g47580<br>• encodes an LRR protein kinase family member |
| 2 | Insertion downstream from At3g03460<br>• encodes an ankyrin repeat family protein |

**Table 2:** Segregation of T-DNA insertions in four target genes in *std1* segregants

| Plant Number | Phenotypic summary | At4g16490 status | At3g21790 status | At3g03456 status | At3g48590 status |
| --- | --- | --- | --- | --- | --- |
| 1 | Short plants with short, thick stems, thick and wavy leaves, altered phyllotaxy, sterile (class 3 plants) | T-DNA | T-DNA | WT | WT |
| 2 |  | WT | T-DNA | WT | WT |
| 10 | Reduced height plants with abnormal phyllotaxy, reduced fecundity (class 2 plants) | WT | T-DNA | WT | WT |
| 14 |  | T-DNA | T-DNA | WT | WT |

**Table 3:** TAIL-PCR of *std2* reveals a single gene target and complex T-DNA integration events

| Sequence count | Identity/annotation |
| --- | --- |
| 14 | T-DNA rearrangement/PCR artifact |
| 10 | Tandem repeat T-DNA |
| 7 | Reads through LB into vector DNA |
| 3 | Gene target (At3g03890) |
| 1 | Converging T-DNA |

### The HOP2 locus is genetically dense

The HOP2pro>HOP2::eGFP reporter construct we used contained a long 5’ region (4kbp) which contains the neighboring gene in the opposite orientation. Only 660bp separates the start codon of this gene, At1g13320, encoding a regulatory subunit of protein phosphatase 2A (PP2A3), and the start codon of HOP2 (Figure 2). At the 3’ end of the reporter gene construct there is a potential overlap in the 3’ regulatory region of a gene in the opposing orientation, At1g13340, encoding ISTL6, which may play a role in regulating multivesicular body formation (Buono et al, 2016). Thus, the presence of parts of these neighboring genes in the transforming construct led us to hypothesize that the T-DNA copy number and/or the sites of insertion may lead to misexpression of any or all of the three genes. We therefore examined the expression of these genes by qRT-PCR (Figure 3). Inflorescence RNA was prepared from L*er* and *std2* plants, and qRT-PCR conducted using control primers for the SAND (At2g28390) gene (Czechowski et al., 2005). We found that HOP2 mRNA levels were over 40 fold higher in *std2* than L*er* (2^ΔΔCT^=5.3), but the levels of PP2A3 and ISTL6 transcripts were also slightly elevated (3 fold and 2.3 fold, respectively).

**Figure 2:**
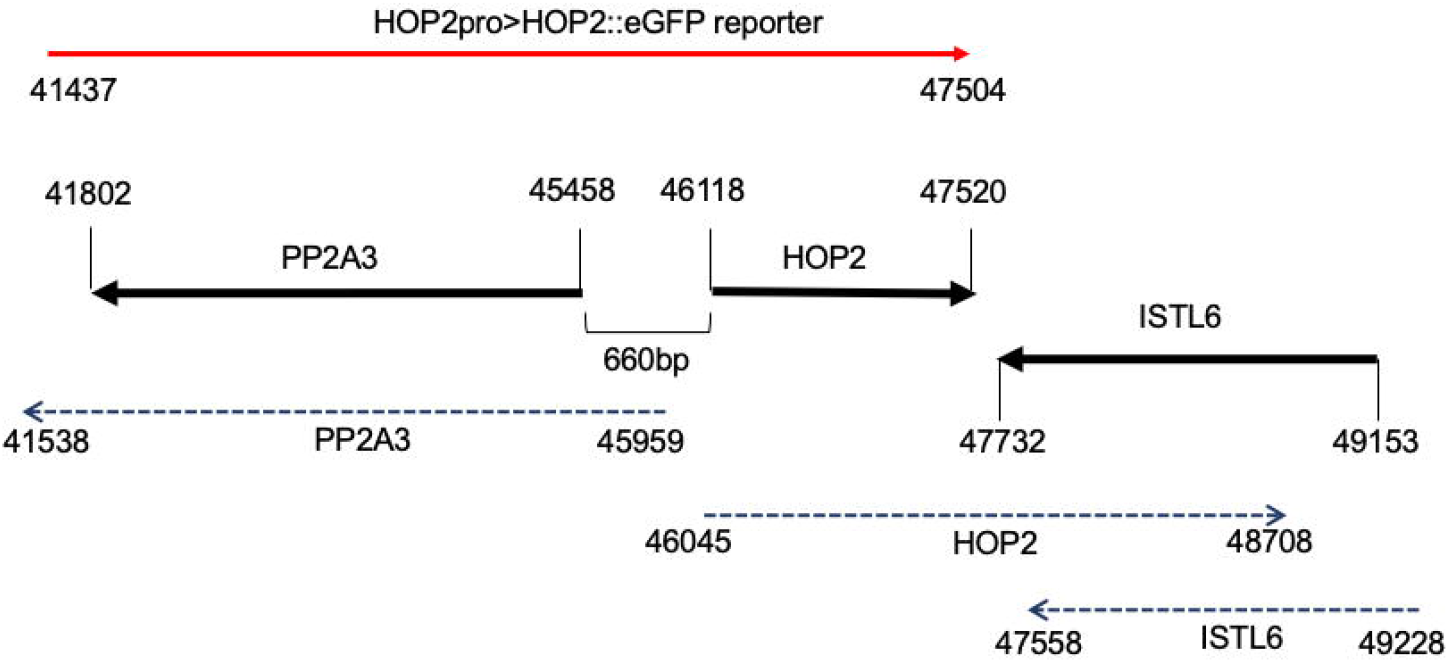
The *HOP2* locus is genetically dense. A schematic of three neighboring and partially overlapping genes: HOP2, PP2A3 and ISTL6, is shown. Numbers refer to the BAC T6J4 coordinates of the start and stop codons of the three genes. The region matching the T-DNA used for transformation is shown in red. Transcript maps are shown by hatched lines with inclusive coordinates.

**Figure 3:**
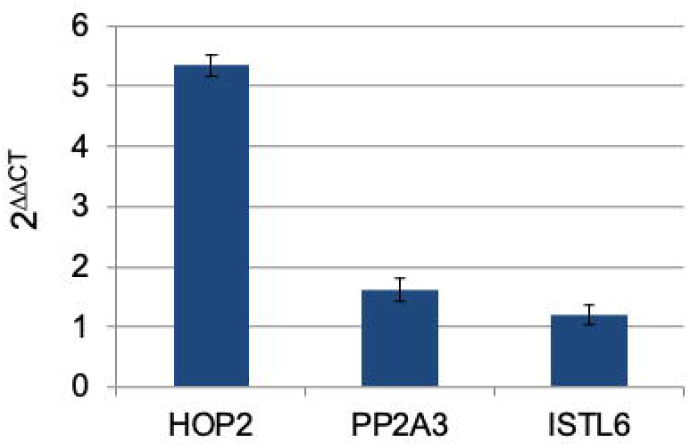
Transcript levels of *HOP2, PP2A3* and *ISTL6* in *std2* inflorescences. Triplicate samples for L*er* and *std1* were subjected to qRT-PCR with primer sets for the three genes. SAND1 was employed as a control primer and 2^ΔΔCT^ values were determined. Error bars represent standard error of the mean.

### Overexpression of HOP2 leads to defects in growth and development

It is conceivable that slight alterations in the expression of PP2A3 and/or ISTL6 could be responsible for the *std* phenotype. To more specifically investigate this issue and to determine if HOP2 overexpression is causal, we generated a dexamethasone inducible construct of HOP2 that has no known PP2A regulatory sequence and ends before the 3’ end of the HOP2 gene, well before any overlap with the converging ISTL6 transcription unit. The construct also encodes an HA tag at the 3’ end. We conducted several experiments to investigate the effect of overexpression of HOP2 on growth and development. First, DEX>HOP::HA (LC842) plants were grown together with wildtype and presented no obvious differences in development when grown on MS medium (Figure 4). However, after transferring seedlings grown on MS medium to medium containing 1μM DEX, DEX>HOP::HA plants remained small and eventually became necrotic and died. Similarly, in soil grown plants, after several foliar applications of DEX to inflorescences, growth of DEX>HOP2::HA plants was compromised (Figure S3) and plants exhibited phyllotaxy defects and reduced fecundity (Figure 5).

**Figure 4:**
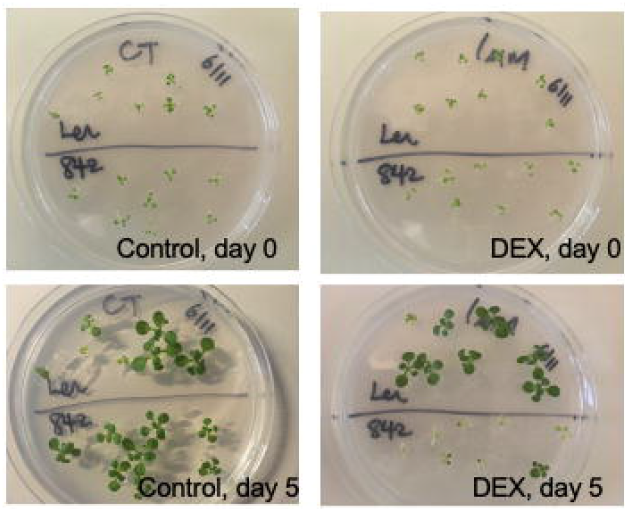
DEX-induced HOP2 compromises plant growth. Seeds of L*er*, and L*er* harboring LC842 (DEX>HOP2::HA) were germinated on MS medium and one week old seedlings were gridded to control (MS) plates, and MS plates containing 1uM DEX. Plates were photographed at day 0 (gridding day) and at day 5 after gridding.

**Figure 5:**
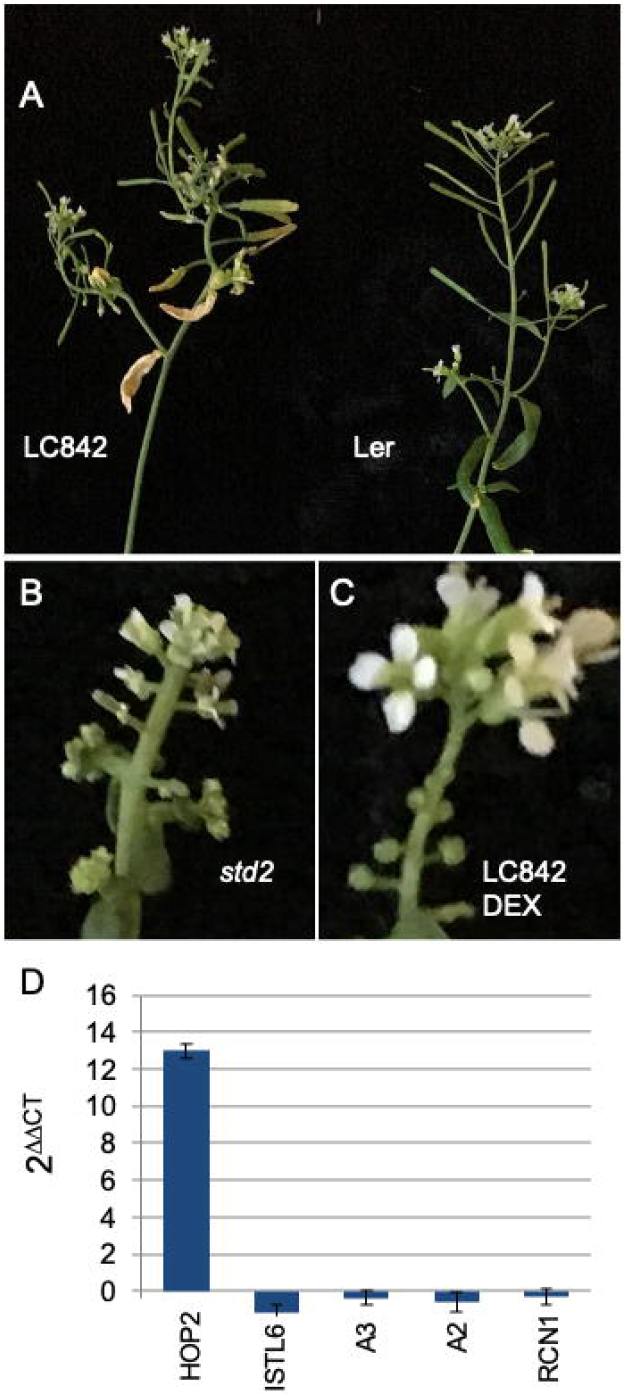
DEX induced HOP2 leads to *std*-like phenotypes. A. Comparison of DEX treated L*er* control plant with LC842. The 842 plants exhibited stunted inflorescence growth, phyllotaxy defects and reduced fecundity. B,C: *std2* and DEX treated LC842 inflorescences, showing similar developmental defects. D. qRT-PCR results of LC842 seedlings grown on either MS medium (control) or 1uM DEX for 3 days. *HOP2* is induced >8200 fold (p=7.4 x 10^-5^), whereas *PP2A3* (A3), *PP2A2* (A2) and *PP2A1* (*RCN1*) show no significant changes in expression (p values >0.093 in all cases). *ISTL6* is downregulated by 2.3 fold (p value= 0.024).

Lastly, we conducted qRT-PCR on seedlings that had been grown on MS media, and then transferred to new MS plates or plates containing 1uM DEX. After 3 days of growth, RNA was prepared and qRT-PCR conducted. We observed that HOP2 was overexpressed by >8200 fold (2^ΔΔCT^=13), whereas there was little change in PP2A3 or ISTL6 levels (Figure 5d). In addition, we measured the expression of two other PP2A family members, PP2A2 and RCN1, which were largely unchanged. Taken together with the phenotypic changes induced by DEX that mimic *std* mutants, these experiments point to elevated levels of HOP2 as causing dramatic changes in growth and development.

## Discussion

Although the basic body plan of *std* mutants is maintained, many aspects of vegetative and reproductive development are affected. Our TAIL-PCR experiments revealed T-DNA insertions near several genes, but these insertions did not faithfully segregate with the *std* mutant phenotype. For three of the five genes, literature reports suggest that there are no obvious phenotypes from T-DNA insertions (At3g21790, Dong et al., 2014; At4g16490, Luhua et al., 2013; At4g34060, Yu et al., 2020), and no phenotypes have been reported for a variety of insertions in the other two genes. These factors led us to further dissect the *HOP2* region, where *PP2AA3* and *ISTL6* genes partially overlap *HOP2*. Our qRT-PCR and DEX-inducible HOP2 experiments strongly suggest that misexpression of neither PP2AA3 nor ISTL6 is responsible for the *std* phenotypes, supporting the conclusion that HOP2 overexpression is causal.

Despite the negative evidence for alterations in PP2A expression as causal, we were intrigued by phenotypes of PP2AA higher order mutants, which are strikingly similar to those of *std* mutants. In particular, *rcn1/pp2a2* and *rcn1/pp2a3* double mutants both compromise plant height and fecundity (Zhou et al., 2004). In addition, the extreme class 3 *std* segregants are phenotypically very similar to the *fass/tonneau2* (*ton2*) mutant in that they exhibit defects in stem elongation and floral organ development, and possess thickened inflorescence stems, thick rumpled leaves, and are sterile (Torres-Ruiz and Jurgens, 1994; Fisher et al., 1996; Camilleri et al., 2002). FASS/TON2 is a regulatory B’’ subunit of PP2A phosphatases (Spinner et al., 2013), that interacts with PP2A1/RCN in the yeast two-hybrid system (Camerilli et al, 2002). Moreover, TON1, a protein essential for preprophase band formation (Traas et al., 1995; Azimzadeh et al., 2008), is also a part of this complex, and the *ton1* mutant phenotype shares similarities with our severe class 3 mutants. Despite the lack of evidence for misexpression of PP2A and TON genes, at least at the transcriptional level, the striking similarities of higher order PP2A and TON mutants with our *std* mutants point to a potential link between HOP2 misexpression and PP2A function. Although PP2A phosphatases control many aspects of cellular metabolism, the misshapen organs and phyllotaxy defects are easily understood in the context of defects in cell division mechanisms and alteration of the planes of division.

How else might HOP2 overexpression underpin the *std* phenotype? In plants, and as a key feature of meiosis, HOP2 plays roles in the pairing of homologous chromosomes, and in promoting interhomolog DNA repair/recombination (Schommer et al., 2003; Vignard et al., 2007; De Muyt et al., 2009; Stronghill, et al., 2010; Uanschou et al. 2013). HOP2 interacts with its partner protein, MND1, to modulate the activity of the RAD51 and DMC1 repair/recombination enzymes (Petukhova et al., 2005; Chi et al., 2007; Bugreev et al., 2014; Zhao et al., 2014; Tsubouchi et al., 2020). Interestingly HOP2 alone can act as a recombinase in vitro (Petukhova et al., 2005). As a protein that fosters recombination, it is conceivable that elevated levels of HOP2 result in somatic recombination, perhaps generating unrepaired templates and/or promoting crossing over between nonhomologous chromosomes that results in chromosome fragmentation and subsequent cell death. Another possibility is suggested by yeast two hybrid experiments in which the rice MND1 protein was shown to interact with replication protein A (RPA; Lu et al., 2020). RPA binds to and stabilizes ssDNA intermediates during DNA replication, repair and homologous recombination. In vitro reconstitution experiments show that HOP2 competes with RPA for binding to ssDNA (Chan et al., 2019). Thus, if HOP2 is overexpressed, ssDNA regions that ordinarily would be stabilized by RPA could assume a conformation that is incompatible with the processes of replication, repair and/or recombination, and failure of these processes (e.g. replication fork collapse) would result in DNA rearrangements and possible cell death. Overexpression of other genes that play roles in DNA/chromosome morphology or behavior has been demonstrated to produce vegetative morphological defects as well as fecundity problems. For example, overexpression of the rice *TOP6A1* gene, the homolog of Arabidopsis *SPO11-1*, produces a variety of developmental defects (Jain et. al., 2008). Similarly, plants overexpressing CTF7ΔB, encoding a kinase that acetylates cohesin complexes and is required for DNA repair, mitosis and meiosis, gives rise to cell division defects as well as a broad range of developmental defects, including dwarfism, fasciation, and phyllotaxy defects (Liu and Makaroff, 2015). It was also recently demonstrated that HOP2 interacts with ATF4, a b-ZIP transcription factor that guides the differentiation of osteoblasts in mice (Zhang et al., 2019). In this regard, Kang and coworkers (2015) conducted molecular dynamics modeling studies of crystallized HOP2/MND1 with DNA, and suggested that the complex distorts the recipient dsDNA. This distortion of the DNA helix could alter the dynamics of transcription factor binding and the overexpression of HOP2 may therefore indirectly alter numerous processes essential for development and physiological homeostasis. Future RNA-seq or microarray analyses could determine whether large scale changes in transcription can be correlated with elevated HOP2 levels.

Cells that have suffered lesions due to faulty repair and/or ectopic recombination events may suffer catastrophic cell death (e.g. by chromosome fragmentation), or may be committed to programmed cell death (PCD). In either case, it is likely that intercellular communication networks are affected, which ultimately would alter hormone homeostasis. In this regard, the release of tryptophan from dying cells is proposed to drive the production of auxin in neighboring cells (Sheldrake and Rupert, 2021), and PCD of root cap cells influences auxin levels and the commitment of adjacent cells into lateral root primordia (Xuan et al., 2016). Some *std* phenotypes are consistent with altered auxin levels and crosstalk between hormonal biosynthesis and response pathways likely initiate a cascade of hormonal adjustments. For example, the defects in gynoecium development we observed for the *std* mutants are reminiscent of the phenotypes of well-known mutants such as *ettin, crabs claw, spatula*, and *hecate*, whose defects are influenced by and/or influence hormone metabolism (reviewed by Marsch-Martinez and de Folter, 2016). In the future, cytological examination of *std* mutants and direct hormone measurements would allow this hypothesis to be tested.

## Acknowledgements

We appreciate the advice on this project and comments on the manuscript by Dr. Clare Hasenkampf (CAH). The authors are grateful to Bruno Chue, Celine Anton, and Ellie Kubisz for their technical assistance. This work was supported by a grant from the Natural Sciences and Engineering Research Council of Canada (NSERC) to CAH and CDR (RGPGP-2015-00071). MK was supported by an NSERC summer fellowship.

