## Supplementary figures and images for "Overexpression of HOP2 induces developmental defects and compromises growth in Arabidopsis"

### Figure S1

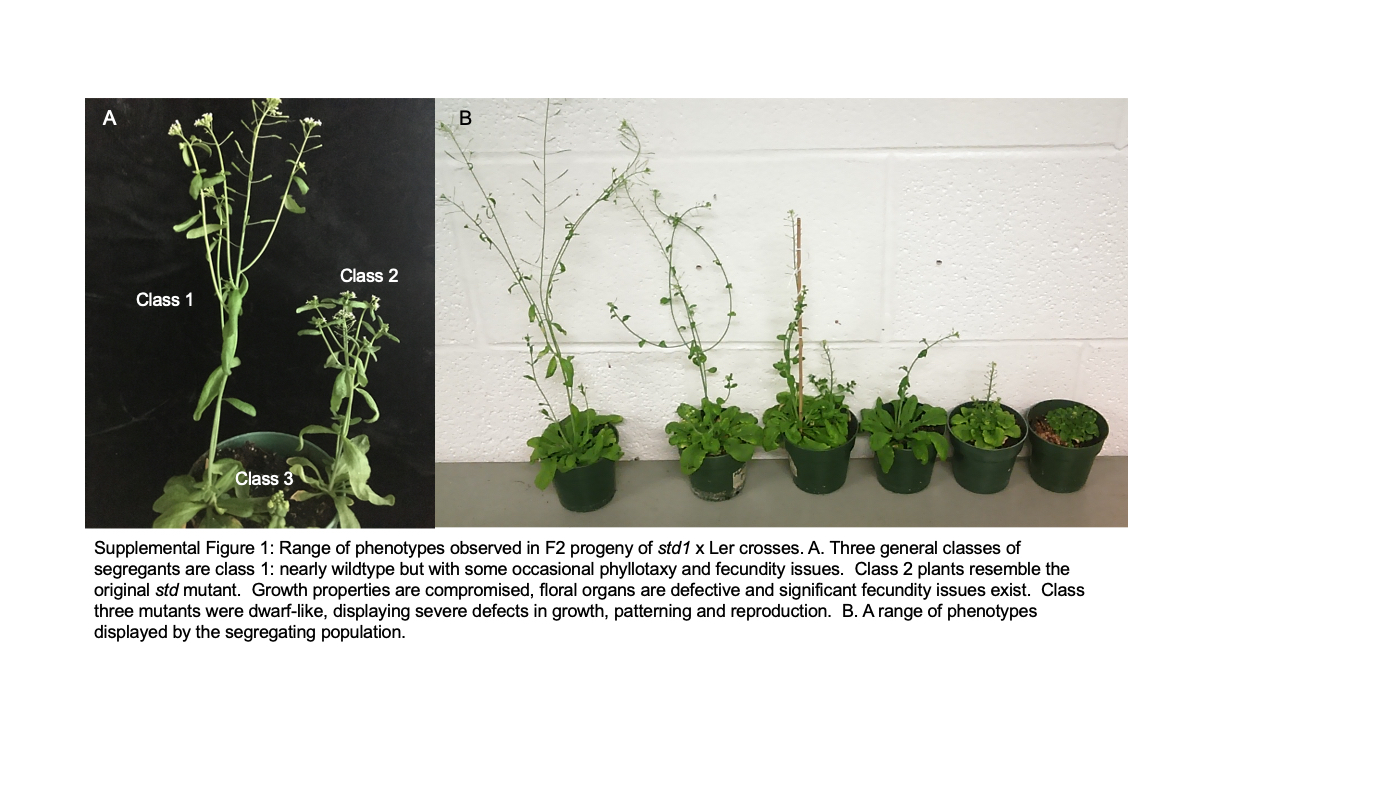

### Figure S2

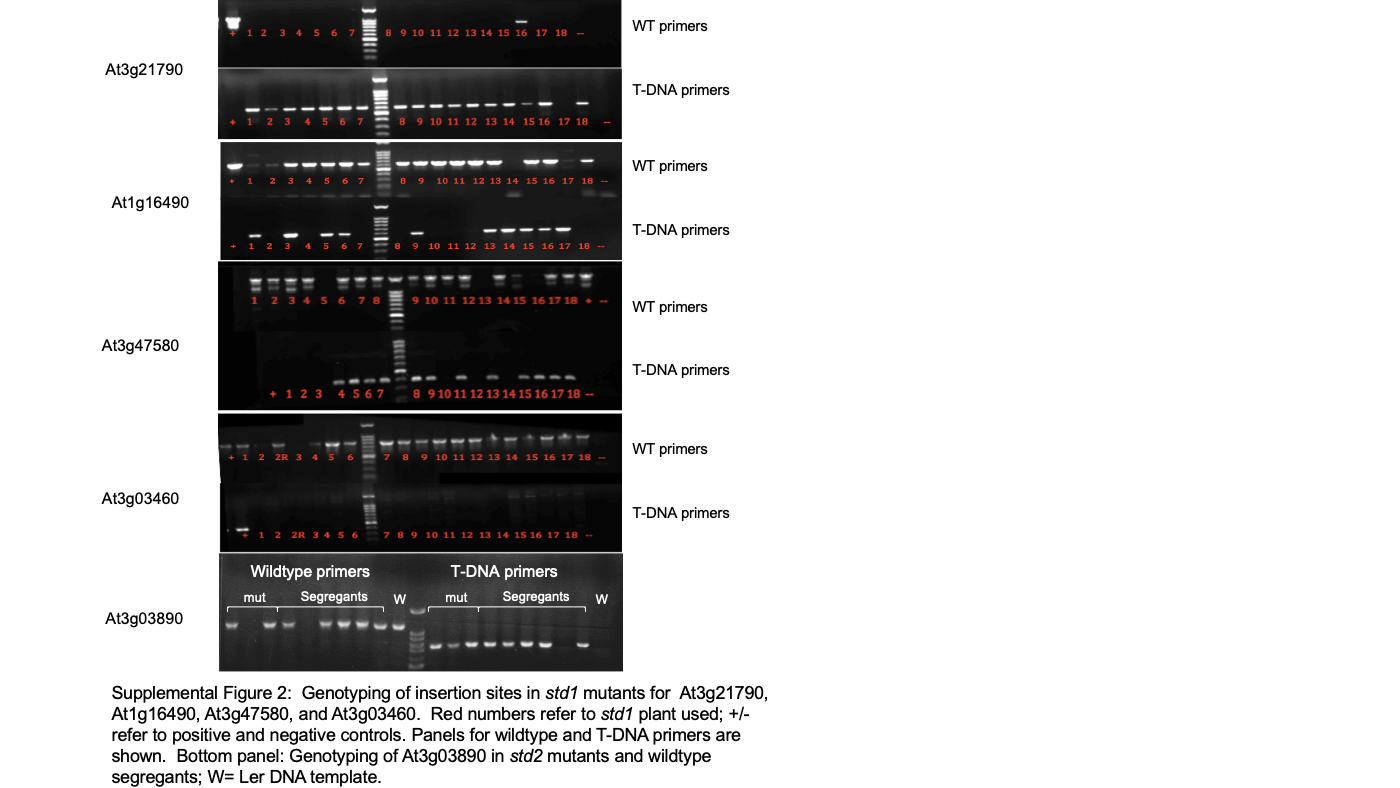

### Figure S3

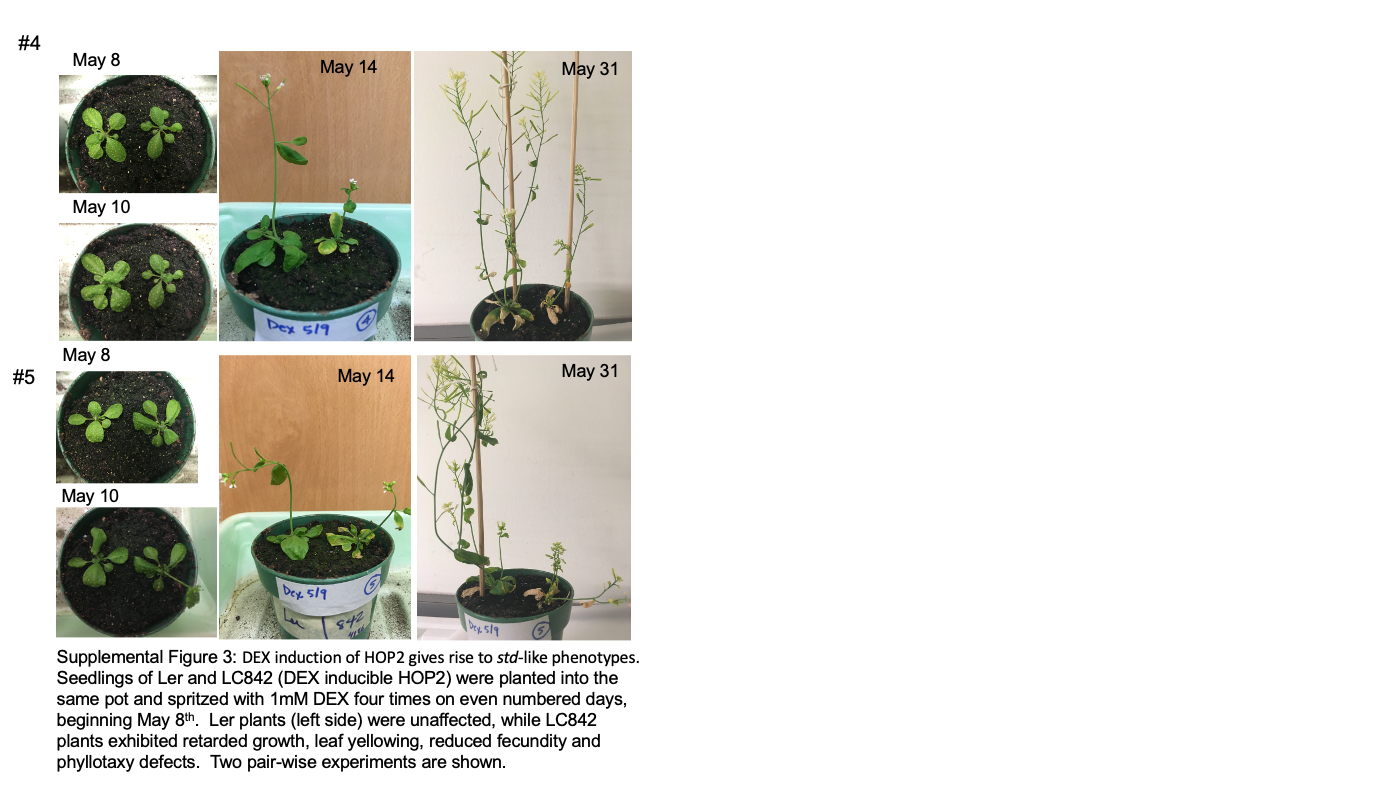
