## Supplementary material for "Overexpression of HOP2 induces developmental defects and compromises growth in Arabidopsis": Table S1

### Table S1: Primers, genotyping data and molecular techniques

Primers for generating HOP2pro>HOP2::eGFP reporter construct

| Primer | Sequence | Amplicon size |
|---|---|---|
| HOP2proFOR Pac | CTTAATTAAAAAGACAACAACGACCTGAATCT | 4692bp |
| HOP2proBACK Asc | TGGCGCGCCCCGATTTAGGAGCCATTTTTGTC | |

These two primers were used to amplify the promoter region to create LC817 having PacI and AscI sites (underlined). This clone is a pEGAD derivative with the coding region of GFP 3’ to the AscI site. Then a FOR6/BACK6 PCR product of 1.7kb was generated, which contains the HOP2 coding region. This fragment was cut with SacI and AscI, and inserted into LC817 which had been cut with the same enzymes, to generate the final HOP2pro>HOP2::eGFP construct.

| Primer | Sequence | Amplicon size |
|---|---|---|
| HOP2 FOR 6 | ATTTACCGATTACGCTGCTTTCCAAGTTCCGAGC | 1711bp |
| HOP2 BACK 6 Asc | ACTGGGCGCGCCGCCTCTTTTTACCATGTT | |

DNA sequencing confirmed the expected in-frame fusion of the HOP2 coding region with that of GFP.

#### Primers for TAIL-pcr

RB gene specific primer T6J4081117: 5’ TACTCCACATATCAGATTCAGGTCGT 3’
LB gene specific primer. LBiPCRFOR042117. 5’ CAAACTGGAACAACACTCAACCCT 3’

Degenerate primers:

AD1 081117- NGTCGASWGANAWGAA
AD2 081117- TGWGNAGSANCASAGA
AD3 081117- AGWGNAGWANCAWAGG
AD6 081117- WGTGNAGWANCANAGA

#### Primers for qRT-PCR

| Primer | Sequence | Amplicon size |
|---|---|---|
| SAND1 FOR | GCGTTAAGGCAAGCTACAGG | 110bp |
| SAND1 BACK | TTTCTGTGCACCAGCAAGAC | |
| PP2A3 FOR | GGAAACTTGCGTGAGGGAGA | 110bp |
| PP2A3 BACK | GCTGAAAGTCGCTTAGCCAG | |
| HOP2 FOR | CCTAAATCGGATAACACCGAAGC | 323bp |
| HOP2 BACK | CTTGATTTCTGATTCCACATCACTG | |
| ISTL6 FOR | AAGATGGCAGAGGAGAAGGT | 190bp |
| ISTL6 BACK | CCCACCAATGTTACCCGGAC | |
| RCN1 FOR | TCCGCACTACCTACACAGGA | 223bp |
| RCN1 BACK | GTCCACCAAACACTGACGGA | |
| PP2A2 FOR | GTGCCTCGGTACGGGTTTTG | 148bp |
| PP2A2 BACK | GGCTTAAGCAGCACGTTGAAAA | |

#### TAIL-pcr targets genotyping primers

| Target gene | Test | Forward primer | Reverse Primer | Amplicon size (bp) |
|---|---|---|---|---|
| At4g16490 | Wild type | GACACTGAAGAGTAAAACCCAAGGC | AACGGAAAGCGAGTGGAATTG | 607 |
| | T-DNA | ATTTTGCCGATTTCGGAAC | AACGGAAAGCGAGTGGAATTG | 617 |
| At3g21790 | Wild type | CCCCACGCTTACGCATCTAAAC | TGCTTCTCACCTTTGGTTCTTGG | 990 |
| | T-DNA | ATTTTGCCGATTTCGGAAC | TGCTTCTCACCTTTGGTTCTTGG | 680 |
| At3g03460 | Wild type | TTTCTTGTAGCTGCGATTGGGAC | CAAACGGCACCAAATGCGTG | 820 |
| | T-DNA | TTTCTTGTAGCTGCGATTGGGAC | ATTTTGCCGATTTCGGAAC | 543 |
| At3g47580 | Wild type | CCTTGTCGTCTTGGTAATTGCG | TTGGCTCACAACCACACAGAATAC | 1527 |
| | T-DNA | ATTTTGCCGATTTCGGAAC | TTGGCTCACAACCACACAGAATAC | 461 |
| At3g03890 | Wild type | CCATTTCATCTTCTGCATCCCTC | TCTGTATCACTGTTGCCACTCCC | 1135 |
| | T-DNA | ATTTTGCCGATTTCGGAAC | CCATTTCATCTTCTGCATCCCTC | 903 |

#### Primers to test T-DNA organization in planta

To test for tandem repeats: EGAD BACK 081117- CCTGACTCCCTTAATTCTCCGCTCAT and LB1.3: ATTTTGCCGATTTCGGAAC were used

To test for T-DNA dimer convergence: newLBA- GCCCTGATAGACGGTTTT was used

To test for T-DNA dimer divergence: EGADback was used

To test for read-through of the left border, the primers newLBA and LC830LB BORDER BACK- CTCATGTTACCGATGCTATTCG were used

#### Primers for Gateway cloning to generate LC842

| Primer | Sequence | Amplicon size |
|---|---|---|
| GATEWAY FOR | GGGGACAAGTTTGTACAAAAAAGCAGGCTCCGACAAAAATGGCTCCTAAATCGG | 1468 |
| GATEWAY BACK | GGGGACCACTTTGTACAAGAAAGCTGGGTCCTGTCCTCGAGGCCTCTTTTTACCAT | |
